# Population density and plant availability interplay to shape browsing intensity by roe deer in a deciduous forest

**DOI:** 10.1101/2021.11.25.468041

**Authors:** William Gaudry, Jean-Michel Gaillard, Sonia Saïd, Anders Mårell, Christophe Baltzinger, Agnès Rocquencourt, Christophe Bonenfant

## Abstract

Browsing damage in forests relies on a complex interaction between herbivore density and both forest understory composition and relative availability. Although variation in the amount of browsed twigs is sometimes used to assess abundance of large herbivores, the potential confounding effect of resource availability on this relationship has not yet been investigated. To fill the gap, we measured how browsing intensity of the woody plants varied in response to changes in both roe deer (*Capreolus capreolus*) abundance and vegetation availability from an intensive long-term monitoring. We estimated plant availability and consumption by roe deer from a modified Aldous method throughout a 14 yearlong period during which we experimentally manipulated population density. The functional response was strongly non-linear and density-dependent. When plant availability was low (< 12.5%), browsing intensity strongly increased with plant availability with an increasing rate with roe deer density, whereas beyond this threshold, browsing intensity slightly increased with both plant availability and population density in an additive way. Thus, forest susceptibility to browsing increases with increasing competition for food, especially when plant availability is low. Moreover, the interplay between browsing intensity and population density at low plant availability prevents the use of browsing intensity to monitor roe deer abundance when plant availability is low. Our findings provide clear evidence that relying on key ecological concepts such as functional responses improves the accuracy of management tools when monitoring changes of the herbivore-plant system over time.

## 1. Introduction

Populations of large herbivores have strongly increased in both range and abundance over the last decades in temperate areas (Côté et al., 2004; Apollonio et al., 2010). This general and continuous increase in the abundance of large herbivores has led to major changes in forest composition (Ramirez et al. 2019). By consuming plant tissues, herbivores affect the growth, survival and reproduction of plants (Hidding et al., 2012; Sabo, 2019). Heavy browsing generally favours the abundance of species resistant to browsing, at the expense of plant species preferred by herbivores (Tremblay et al., 2006; Hidding et al., 2013; Perea et al., 2014). In addition, the high browsing pressure exerted by abundant herbivores has strongly affected the forest landscape, mainly by reducing availability of understory (Fuller, 2001; Rooney and Waller, 2003). Overall, by affecting tree recruitment and forest regeneration, browsing by ungulates is currently considered as one of the main threats to forest (Schuck and Requardt, 2008; Perea et al., 2014; Ramirez et al. 2019).

The ecological consequences of browsing on plants depend on both deer selectivity and on the tolerance of plants to browsing (Long et al., 2007; Perea et al., 2014; Sabo, 2019). Availability of understory vegetation is known to be one of the key factors affecting the sensibility of forest regeneration to browsing damages (Gill, 1992; Vospernik and Reimoser, 2008). The amount of understory vegetation available to large herbivores is closely linked to the dynamics of overstory vegetation (Gill et al., 1996; Vospernik and Reimoser, 2008). In most exploited forests, the removal of canopy increases the amount of understory vegetation available to herbivores following man-made or natural clearing (Gill et al., 1996; Olesen and Madsen, 2008; Vospernik and Reimoser, 2008). As the forest stand grows up, canopy closure leads the amount of understory vegetation available for herbivores to decrease drastically (Gill et al., 1996; Olesen and Madsen, 2008; Vospernik and Reimoser, 2008). To understand better the forest-large herbivore system and take the appropriate management rules, we need to decipher how variation in population abundance and plant availability shapes browsing intensity (Côté et al., 2004; Tremblay et al., 2006). Relying on the concept of functional response (Holling, 1959), plant consumption should increase non-proportionally with plant availability (Abrams, 1982; Spalinger and Hobbs, 1992), leading intensity of browsing to increase faster at low plant availability and to saturate at high plant availability (type II functional response). Moreover, for a given plant availability, browsing intensity should obviously increase with population abundance. While the latter relationship has been intensively investigated for a long time (see e.g. Aldous, 1944; Morellet et al., 2001; Chevrier et al., 2012), the expected confounding effect of functional response has been mostly overlooked in studies of plant-herbivore systems at very fine spatial scale (i.e at the scale of plant consumption, 4^th^ order selection Johnson, 1980). We aim to fill this knowledge gap by measuring browsing intensity in relation to both abundance of large herbivores and plant availability. We took benefit from an experimental manipulation of roe deer abundance in the intensively monitored forest population of Trois-Fontaines, France, in which browsing intensity and plant cover were recorded over 14 years following a slightly modified version of the method developed by Aldous (1944) and adapted by Ballon et al., (1992). We expected (1) browsing intensity to increase with roe deer density (Morellet et al., 2001; Tremblay et al., 2006; Chevrier et al., 2012), (2) a functional response to occur so that browsing intensity should not increase proportionally with plant availability (Illius et al., 2002; Van beest et al., 2016), and (3) the functional response to be density-dependent, with increasing roe deer density leading to increase intraspecific competition for food (Gill et al., 1996).

## 2. Materials and methods

### 2.1. Study site

We conducted our study in the Territoire d’Étude et d’Expérimentation of Trois-Fontaines, a 1,360 ha enclosed forest located in North-East France (Champagne-Ardennes, 48°43’ N, 2°61’ E) managed by the National Forest Office (Figure 1). The climate is quite mild in winter (mean daily temperature of 2° C in January) and warm in summer (mean daily temperature of 19° C in July), which is favourable to plant growth. Most precipitation comes as rain and rainfall is evenly distributed throughout the year. The soil is fertile and the forest highly productive as indicated by the long-term average of 5.92 m3 of wood produced/ha per year (Inventaire Forestier National). The forest overstory is dominated by oak (*Quercus spp*.; 63.5% of timber trees) and beech (*Fagus sylvatica*, 19% of timber trees) and the understory by hornbeam (*Carpinus betulus*), ivy (*Hedera helix*) and brambles (*Rubus sp*.), which are preferred food resources by roe deer (Duncan et al., 1998). Indeed, the study site offers rich habitat highly favourable for roe deer (Gaillard et al., 1993; Gaudry et al. 2018).

**Figure 1:**
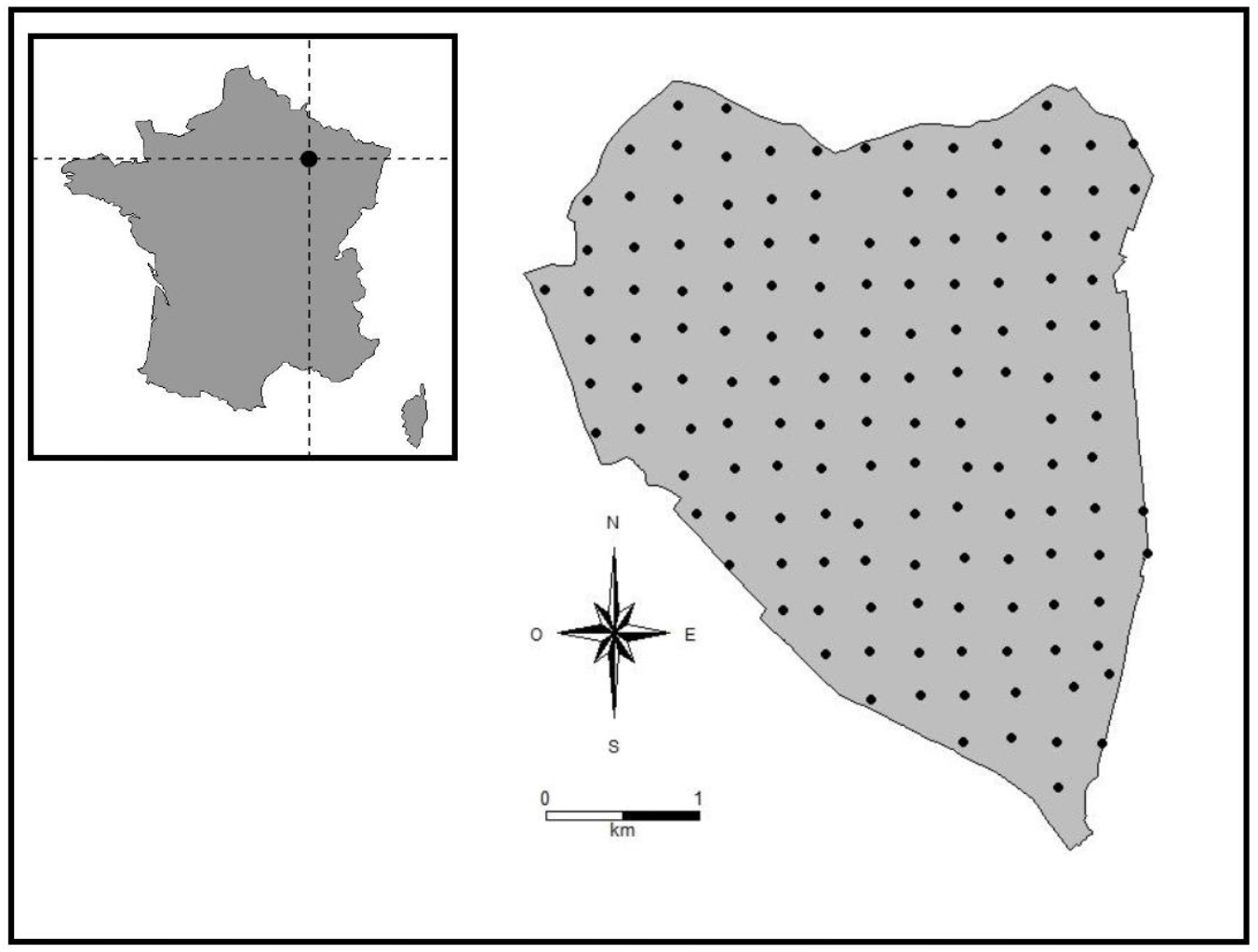
Maps of the Territoire d’Étude et d’Expérimentation of Trois-Fontaines with its location in France. Black points represent the 144 Aldous sampling plots. We monitored plant availability and browsing intensity by roe deer (Capreolus capreolus) in each plot from 1996 to 2009 but in 2000, 2006 and 2008 because of logistic issues.

### 2.2 Roe deer population

The roe deer population is intensively monitored since 1975 using a capture-recapture design (e.g. Gaillard et al., 1993, 2003). From 1977 to 2000, population density was quite constant at about 14.7-18.4 roe deer/km^2^ (200-250 individuals > 1 year of age) thanks to removals during captures (Gaillard et al., 2003). During this period, the demographic performance of the roe deer population was high with virtually all 2-years-old females giving birth (Gaillard et al., 1998) and an annual survival rate of 0.85 and 0.93 for adult males and females, respectively (Gaillard et al., 1993). Then, from 2000 to 2005, roe deer removals stopped as part of an experimental manipulation of density (Chevrier et al., 2012). As a result, roe deer abundance increased markedly and peaked in 2005 at about 32.6 roe deer/km^2^ (mean=443 [95% CI: 360-565] individuals; Figure 2, Gaillard et al., 2013), when roe deer removals started again. This experimental manipulation of density led to density-dependent responses in fawn survival and body mass (Unpubl. Data) and an estimated 15% decrease in average population growth rate (Nilsen et al., 2009). This clear evidence of density-dependence in individual life history traits and population growth demonstrates that intraspecific competition for food among individuals was high right after the density manipulation.

**Figure 2:**
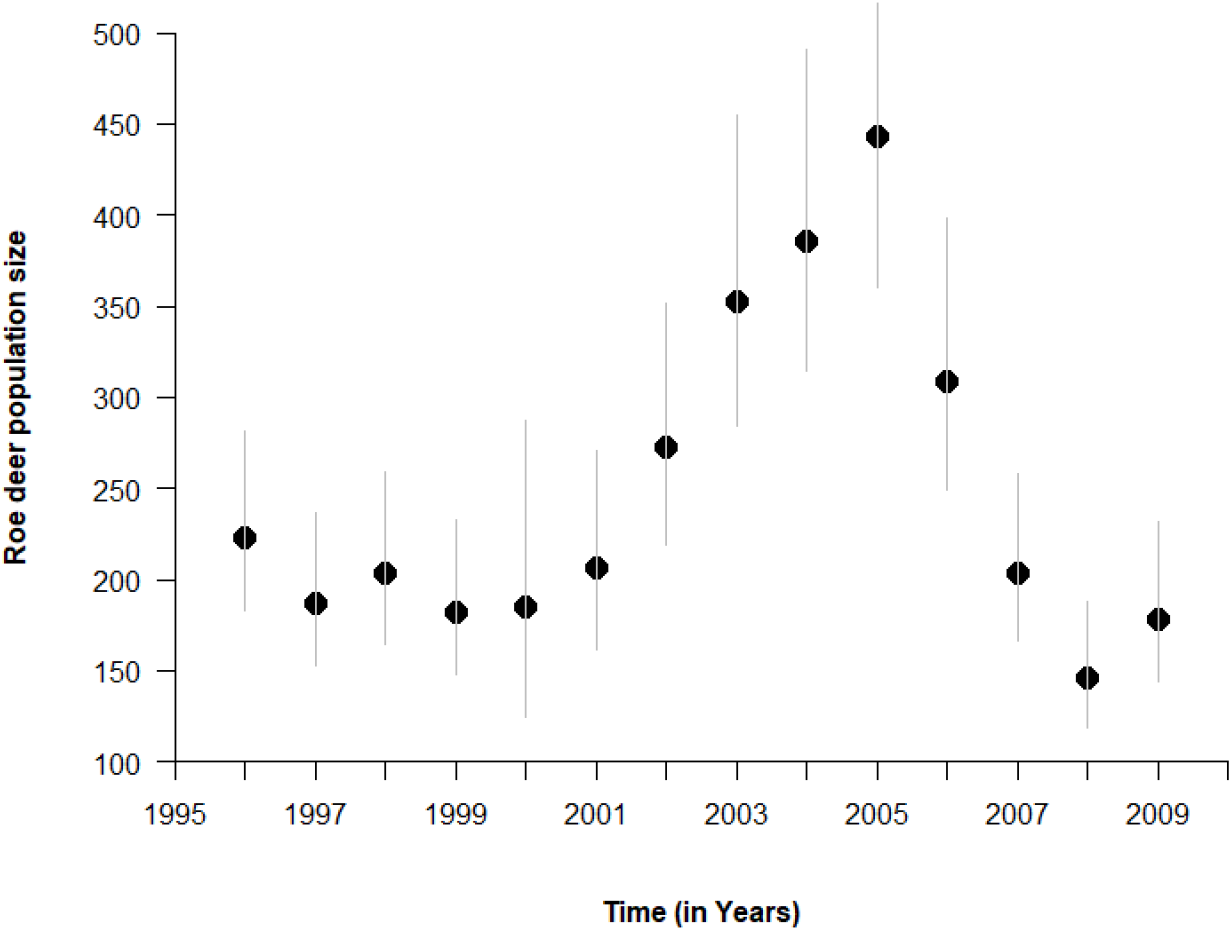
Yearly variation in population size of roe deer (in March) during the study period (1996-2009) at the Territoire d’Étude et d’Expérimentation of Trois-Fontaines (France). Population size was estimated as the number of roe deer older than 1 year of age using capture-recapture methods (see e.g. Gaillard et al., 2013). Vertical gray lines represent the 95% confidence interval of roe deer population size estimates.

### 2.3 Plant sampling

We monitored food resource use and availability by roe deer according to a slightly modified version of the method developed by Aldous (1944) to assess deer impact on vegetation (Ballon et al., 1992). We carried out vegetation surveys between 1996 and 2009 except in 2000, 2006 and 2008 (*t* = 11 years). To obtain a measure of plant consumption over the winter, vegetation surveys took place in the following March-April, before the start of vegetation growth in spring (Morellet et al., 2001). We sampled vegetation at each corner of a grid with a 300×300 m unit cell (1plot/9ha; Figure 1). Overall, we sampled 144 vegetation plots of 40 m^2^. For each sampling plot (that were both geo-referenced by ground marking and geo-located using GPS to allow repeated sampling over years), we exclusively considered woody and semi woody plants potentially available to roe deer (i.e. < 120 cm above the ground). We then measured resource availability and use for each woody and semi-woody plant species detected in the sampling plot. We measured resource availability (hereafter “plant availability”) as the proportion of the sampling plot area covered by each detected plant species (plant cover percentage). Resource use (hereafter “browsing intensity”) was measured as the proportion of buds and shoots browsed for a given plant species.We recorded all plants with presence of scars on the last year’s growth (shoots of the growing season preceding the inventory) resulting from roe deer consumption. The proportion of browsed individual plant defined the browsing index *b*, our response variable. We excluded ivy (*Hedera helix*) from our analyses because consumption is especially difficult to assess in this species (Morellet et al., 2001).

Following Aldous (1944), we classified abundance and browsing intensity into 6 categories: absence (for browsing intensity only because when a given species was not present, it could not be consumed); < 1%; 1% to 5%; 5% to 20%; 20% to 50%; 50% to 75% and >75%. We only considered species that were present in at least 10 % of the sampling plots every year to get accurate estimates of the effect of plant availability and roe deer density on browsing intensity (Morellet et al., 2001). Those measurements assumed that plant availability (*i.e*. plant cover) and browsing intensity (i.e. the proportion of buds and shoots consumed) were good proxies for food availability and food intake by roe deer, respectively. Our dataset then consisted into repeated measurements at each sampling plot, the number of repetition being the diversity of woody and semi-woordy plant species present on the locations.

### 2.4. Data analyses

We analysed the temporal variation of browsing intensity with Linear Mixed Models (LMM, e.g. Bolker et al., 2009). Being a proportion of browsed plants bounded between 0 and 1, we transformed the response variable with a logit function and then tested for the influence of both roe deer density and plant availability. We included the sampling plot identity (N = 144) as a random factor in all models to account for pseudo-replication (*sensu* Hurlbert, 1984). We fitted different candidate models to assess how browsing intensity varied in relation to both roe deer density and plant availability, the latter effect corresponding to a functional response when not proportional (Table 1, see Dupke et al. 2021 for functional response in the context habitat selection). We tested for both additive and interactive effects of these two independent variables to assess whether the relationship between plant availability and browsing intensity changed with roe deer density. We also investigated the shape of the functional response by fitting second-order polynomial and threshold models of plant availability. In threshold models, the interactive effect between density and plant availability can occur either before or after the threshold value of plant availability. We selected models using the Akaike Information Criterion (AIC) as recommended by Burnham and Anderson, (2002). To do so, we fitted mixed models using the maximum likelihood method to conduct model selection but reported parameter estimated from models fitted using the restricted maximum likelihood (REML, Pinheiro and Bates, 2000). We used the *lme4* package (Bates et al., 2011) for R (R Version 3.6.3, www.r-project.org) to fit the LMMs to our data.

**Table 1:**
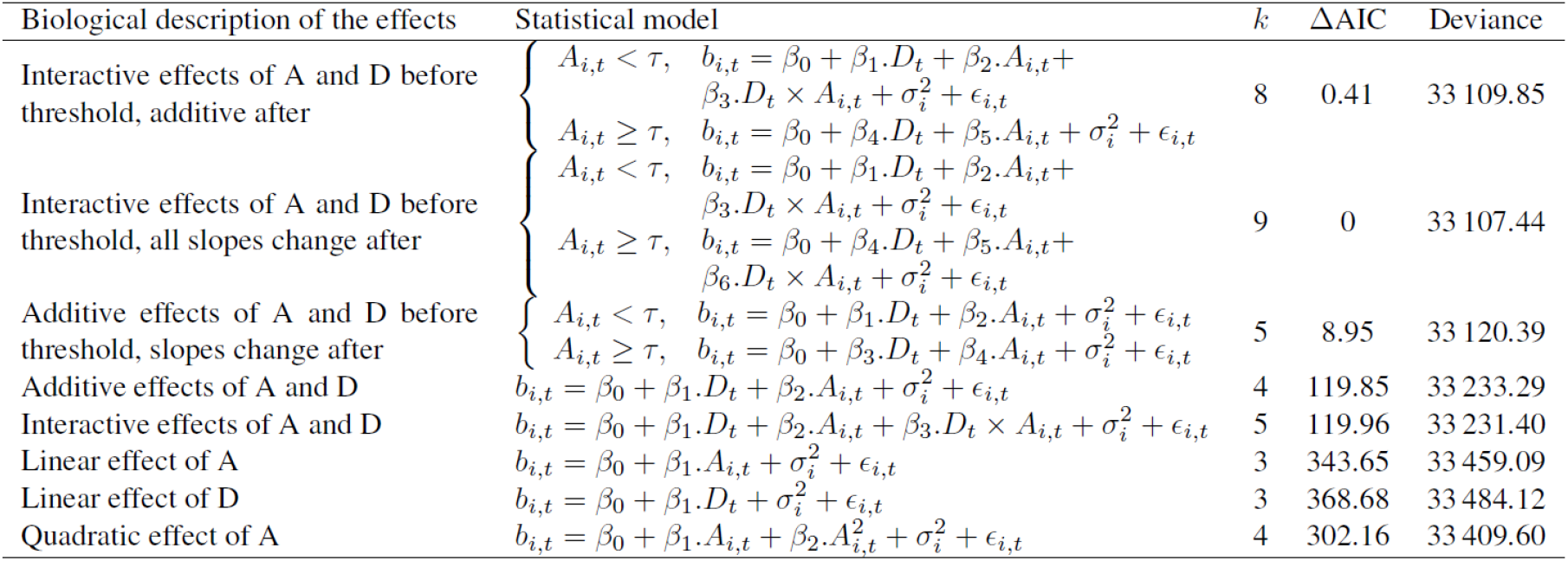
Candidate models (Linear Mixed Models) fitted to investigate the response of browsing intensity to changes in roe deer density (D) and plant availability (A) in the Territoire d’Étude et d’Expérimentation of Trois-Fontaines (France) between 1996 and 2009. Browsing intensity (b_i,t_) is expressed as the percentage of the number of buds and shoots browsed on a logit-scale for the sample plots _i_ at the time _t_. Sample plot identity (N = 144 sample plots) is included as a random factor (σ). β_x_ corresponds to model coefficient and ϵ corresponds to the error term. Candidate models are ranked according to the Akaike Information Criterion (AIC). k represents the number of parameters of the focal model, ΔAIC is the difference in AIC between the candidate model and the model with the lowest AIC. We selected the most parsimonious model among models with a ΔAIC of 2.

| Biological description of the effects | Statistical model | $k$ | $\Delta AIC$ | Deviance |
| --- | --- | --- | --- | --- |
| Interactive effects of A and D before threshold, additive after | $\begin{cases} A_{i,t} < \tau, & b_{i,t} = \beta_0 + \beta_1.D_t + \beta_2.A_{i,t} + \beta_3.D_t \times A_{i,t} + \sigma_i^2 + \epsilon_{i,t} \\ A_{i,t} \geq \tau, & b_{i,t} = \beta_0 + \beta_4.D_t + \beta_5.A_{i,t} + \sigma_i^2 + \epsilon_{i,t} \end{cases}$ | 8 | 0.41 | 33 109.85 |
| Interactive effects of A and D before threshold, all slopes change after | $\begin{cases} A_{i,t} < \tau, & b_{i,t} = \beta_0 + \beta_1.D_t + \beta_2.A_{i,t} + \beta_3.D_t \times A_{i,t} + \sigma_i^2 + \epsilon_{i,t} \\ A_{i,t} \geq \tau, & b_{i,t} = \beta_0 + \beta_4.D_t + \beta_5.A_{i,t} + \beta_6.D_t \times A_{i,t} + \sigma_i^2 + \epsilon_{i,t} \end{cases}$ | 9 | 0 | 33 107.44 |
| Additive effects of A and D before threshold, slopes change after | $\begin{cases} A_{i,t} < \tau, & b_{i,t} = \beta_0 + \beta_1.D_t + \beta_2.A_{i,t} + \sigma_i^2 + \epsilon_{i,t} \\ A_{i,t} \geq \tau, & b_{i,t} = \beta_0 + \beta_3.D_t + \beta_4.A_{i,t} + \sigma_i^2 + \epsilon_{i,t} \end{cases}$ | 5 | 8.95 | 33 120.39 |
| Additive effects of A and D | $b_{i,t} = \beta_0 + \beta_1.D_t + \beta_2.A_{i,t} + \sigma_i^2 + \epsilon_{i,t}$ | 4 | 119.85 | 33 233.29 |
| Interactive effects of A and D | $b_{i,t} = \beta_0 + \beta_1.D_t + \beta_2.A_{i,t} + \beta_3.D_t \times A_{i,t} + \sigma_i^2 + \epsilon_{i,t}$ | 5 | 119.96 | 33 231.40 |
| Linear effect of A | $b_{i,t} = \beta_0 + \beta_1.A_{i,t} + \sigma_i^2 + \epsilon_{i,t}$ | 3 | 343.65 | 33 459.09 |
| Linear effect of D | $b_{i,t} = \beta_0 + \beta_1.D_t + \sigma_i^2 + \epsilon_{i,t}$ | 3 | 368.68 | 33 484.12 |
| Quadratic effect of A | $b_{i,t} = \beta_0 + \beta_1.A_{i,t} + \beta_2.A_{i,t}^2 + \sigma_i^2 + \epsilon_{i,t}$ | 4 | 302.16 | 33 409.60 |

## 3. Results

The observed plant availability was smaller than 12.5% for 90% of our measurements (Figure 3). Two models provided similar fit (ΔAIC = 0.41) and were better than any other one (Table 1). We retained the model with less parameters to comply with parsimony rules. As expected, the relationship between browsing intensity and plant availability was not linear (Figure 4), meaning that a functional response did occur. The model best describing our data included a plant availability threshold of 12.5 %. When plant availability was low (i.e. <12.5%), browsing intensity strongly increased with plant availability, whereas beyond this threshold, browsing intensity slightly increased with plant availability. Moreover, as expected, the functional response was density-dependent. When plant availability was low (i.e. < 12.5%), the increase of browsing intensity with plant availability was faster with increasing roe deer density. On the other hand, at moderate or high plant availability (i.e. > 12.5%) roe deer density and plant availability had additive effects on browsing intensity (Figure 4; Table 2). As expected, for a given plant availability, a greater proportion of plants were browsed at higher roe deer population density.

**Figure 3:**
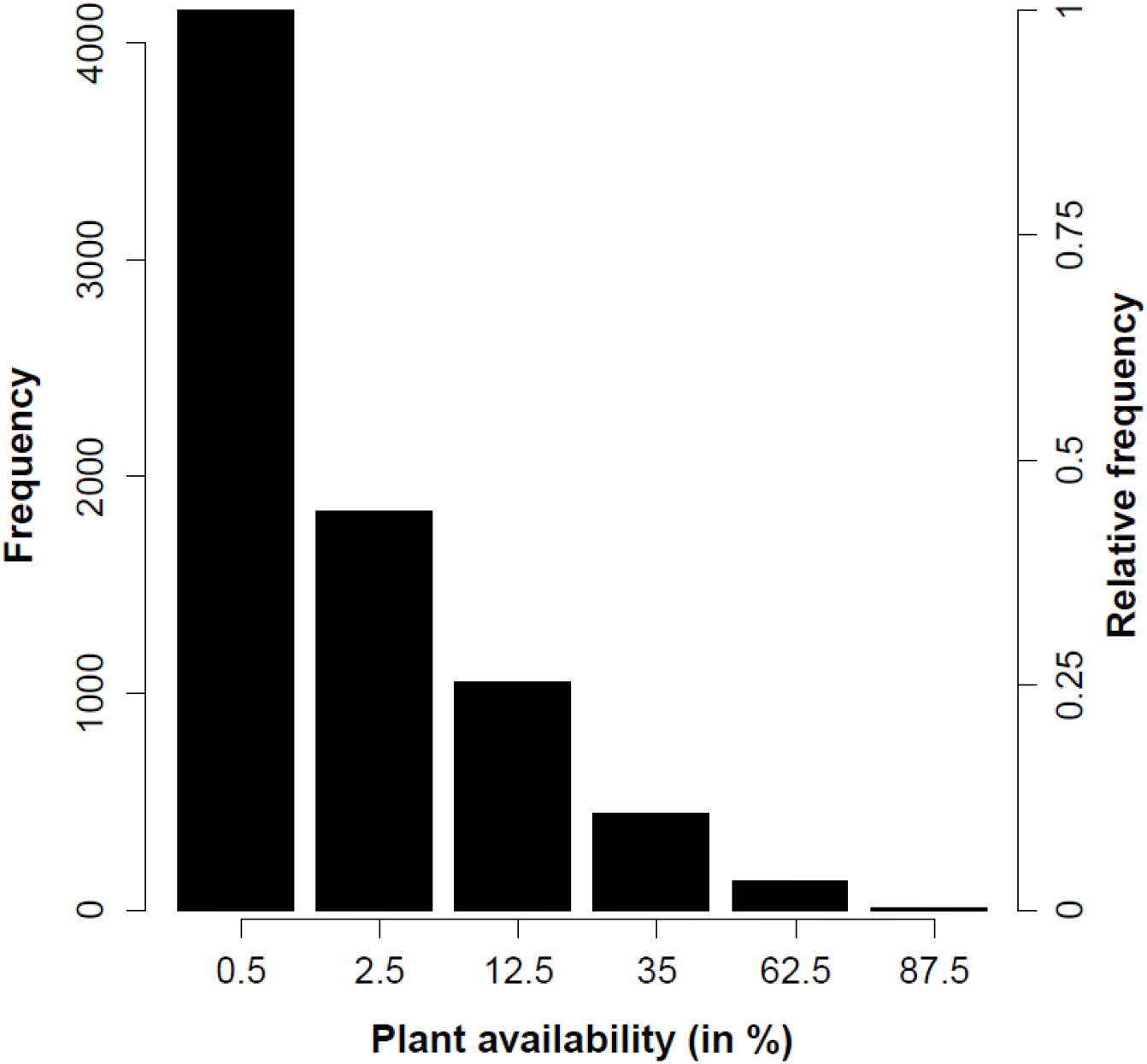
Frequency distribution of plant availability per species measured during the 14-yearlong (1996-2009) vegetation surveys on the 144 sampling plots in the Territoire d’Étude et d’Expérimentation of Trois-Fontaines (France).

**Figure 4:**
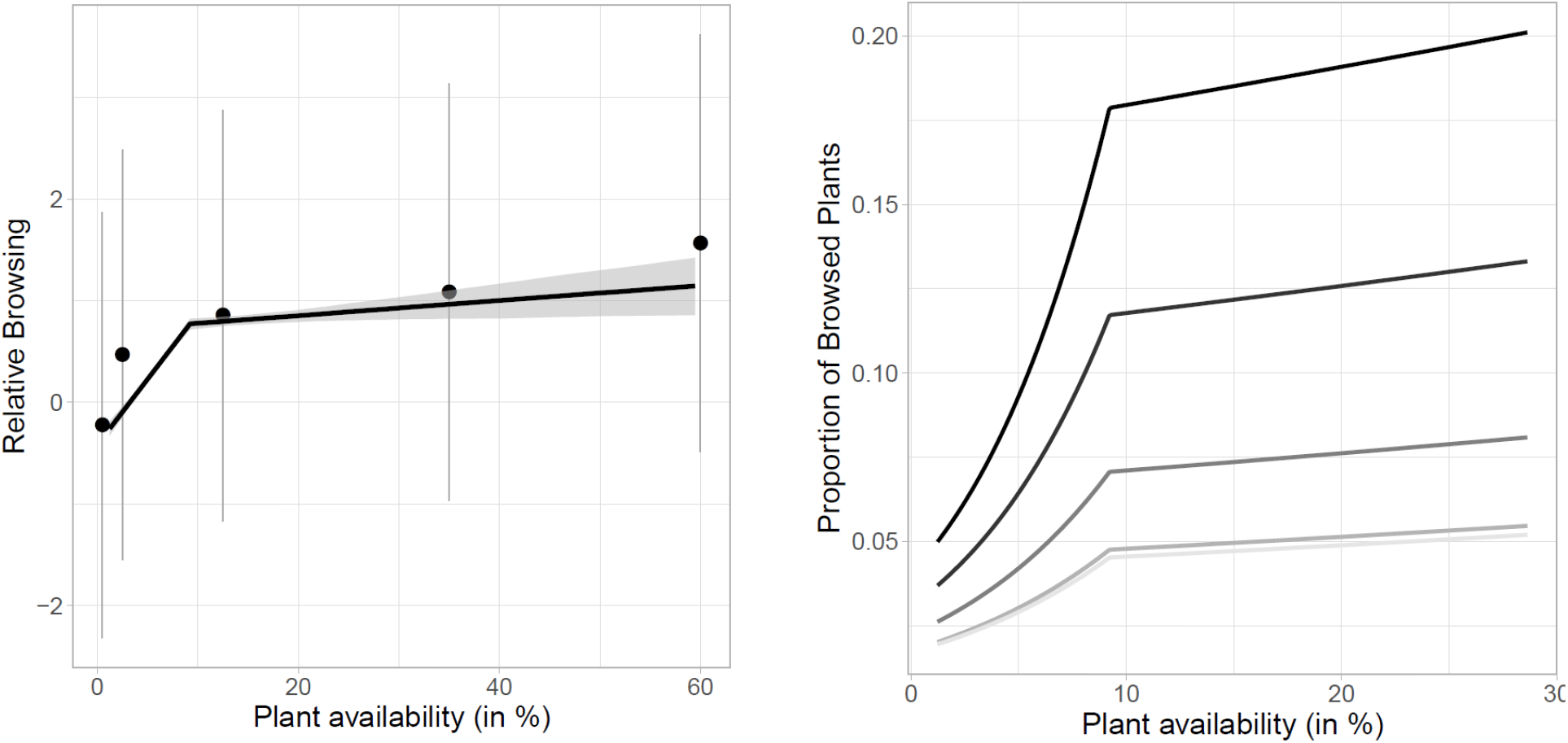
Change in browsing intensity of woody plants by roe deer as a function of woody plant availability in the Territoire d’Étude et d’Expérimentation of Trois-Fontaines (France) between 1996 and 2009 (left); change in proportion of browsed plants in relation to plant availability for different roe deer density. The lowest grey line represents the minimum roe deer density we observed in our study site (10.7 roe deer/km^2^) whereas the upper black line represents the maximum roe deer density (32.6 roe deer/km^2^. Intermediate lines represent the first quartile (13.6 roe deer/km^2^), the median (15.0 roe deer/km^2^) and the third quartile (22.1 roe deer/km^2^) of the estimated annual roe deer density.

**Table 2:**
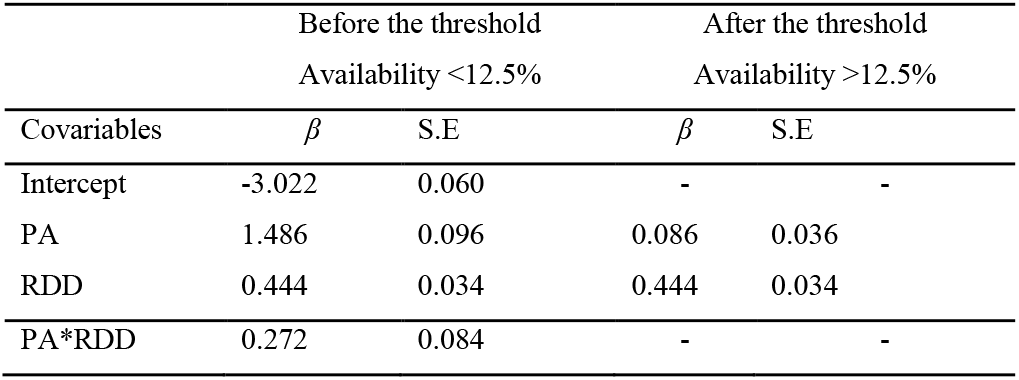
Effect of roe deer density “RDD”, plant availability “PA “ on the browsing intensity on woody plants by roe deer (measured as the percentage of number of buds and shoots browsed) in the Territoire d’Étude et d’Expérimentation of Trois-Fontaines (France) between 1996 and 2009. For each covariate and its first order interaction, we report the estimated coefficient (β) and associated standard error (SE) returned by the mixed model. We entered the vegetation sample plot identity (N = 144 plots) as a random factor in the model (Random effect variance = 0.306; SD = 0.553).

| Covariables | Before the threshold<br>Availability <12.5% |  | After the threshold<br>Availability >12.5% |  |
| --- | --- | --- | --- | --- |
| | $\beta$ | S.E | $\beta$ | S.E |
| Intercept | -3.022 | 0.060 | - | - |
| PA | 1.486 | 0.096 | 0.086 | 0.036 |
| RDD | 0.444 | 0.034 | 0.444 | 0.034 |
| PA*RDD | 0.272 | 0.084 | - | - |

As our sampling design of vegetation plots is a 300×300m grid overlaid on the study area, spatial-autocorrelation might occur in our data, breaking the assumption of independence of observations of generalized linear models (GLM). However, despite the evidence for a spatial structure in our data, accounting or not for spatial auto-correlation did not influence our best model estimates (see appendix 1).

## 4. Discussion

As expected from the concept of functional response, the browsing intensity of roe deer increased with plant availability following a non-linear relationship. This differential use of resource in relation to its availability has frequently been reported in the context of habitat selection at both large (Mysterud ans Ims, 1998; Van Beest et al., 2016; Gaudry et al., 2018) and fine (Shipley and Spalinger, 1995; Illius et al., 2002; Hobbs et al., 2003) spatial scales. Such relation has also been reported in the context of food acquisition as originally proposed by Holling (1959) in predator-prey systems, but at the very fine scale of individual foraging of large herbivores (e.g. Spalinger and Hobbs, 1992). We provide the first clear evidence that a functional response also occurs in browsing intensity at the population scale, which has marked consequences for population management. The rate of increase in browsing intensity by roe deer with increasing plant availability was much higher at low (<12.5%) than moderate or high (>12.5%) plant availability. Moreover, the slope of the functional response of browsing intensity increased with roe deer population density at low plant availability, whereas it remained unchanged at moderate or high plant availability, where increasing density led the browsing intensity to increase by the same amount whatever the exact value of plant availability. At low plant availability, browsers like roe deer should prioritize food quantity over food quality by consuming a high proportion of the available resource, as previously reported in experimental conditions for browsers (Vivas and Sæther, 1987; Shipley and Spalinger, 1995). At moderate or high plant availability browsers select the best quality resource (Illius et al., 2002) and eat a smaller proportion of plants in the high quality food patches (Vivas and Sæther, 1987).

Functional response of roe deer browsing to changes of plant availability was density-dependent at low plant availability. While habitat selection is known to be a density-dependent process (Blix et al. 2014, Van Beest et al., 2016), few studies have yet reported a density-dependent functional response at the scale of the feeding patch in free-ranging large herbivores (but see Blix et al. 2014 on domestic sheep, *Ovis aries*). Previous studies that demonstrated a link between browsing pressure and density of large herbivores did not account for variation in plant availability (Morellet et al., 2001; Chevrier et al., 2012). We provide a clear demonstration that the relationship linking browsing intensity and roe deer density varies in relation to plant availability. When plant availability is low and roe deer density increases, competition for food intensifies to a point where roe deer become less selective and consume a higher proportion of the available food resources (Gill et al., 1996; Molvar and Bowyer, 1994). Similar behavioural adjustment of resource selection by large herbivores has also been reported for captive fallow deer (*Dama dama*) in a manipulative experiment (Bergvall et al., 2006). Likewise, at large spatial scales, both sheep and moose decrease selectivity of productive habitats with increasing interspecific competition for food (Blix et al., 2014; Van Beest et al., 2016). A decreasing selectivity when competition increases seems thus to be a general patterns among large herbivores. When plant availability becomes less limiting, no more behavioural shifts occur in roe deer browsing and individuals of this highly selective forager consistently prioritize food quality over food biomass (Illius et al., 2002; Gaudry et al., 2018).

These findings have important consequences for management of the forest-large herbivore system. They indicate that forest regeneration is less sensitive to browsing by large herbivores at high or moderate food availability (Reimoser and Gossow, 1996). On the other hand, poor forest stands where little or no forest regeneration takes place are exposed to severe browsing damages and increasing roe deer density is expected to have a multiplicative effect on browsing damages (Gill et al., 1996; Vospernik and Reimoser, 2008). Likewise, our findings provide a mechanistic understanding for the lower browsing intensity when increasing the availability of alternative food resources, as previously reported in large herbivores in temperate forests. (Reimoser and Gossow, 1996; Partl et al., 2002; Vospernik and Reimoser, 2008).

Considering the increase in ungulate-human conflicts, tracking changes of the large herbivore-forest system over time is one of the most important challenges that both wildlife and forest managers are currently facing (Coté et al., 2004; Mysterud, 2006). Our analysis of the response of browsing intensity to changes of population density of herbivores (deer density-related response) and plant availability (functional response) in an intensively monitored population of roe deer revealed a complex interplay among those variables. The interplay between plant availability and population abundance we point out in this work prevents a direct and simplistic interpretation of changes in browsing intensity in terms of changes in the abundance of herbivore population. In particular, when plant availability is low, browsing intensity does not display a simple response to population density and should be interpreted with caution in terms of roe deer abundance. On the other hand, when plant availability is moderate or high, the value of the browsing intensity at a given plant availability can be reliably interpreted in terms of roe deer abundance (sensu Morellet et a. 2001). Note, however, that temporal variation in plant availability is likely to occur in response to either natural forest succession stage (Gill et al., 1996; Partl et al., 2002) or forest management and exploitation (Reimoser and Gossow, 1996; Vospernik and Reimoser, 2008). Therefore, the existence of a complex functional response of roe deer browsing requires management tools based on browsing intensity to account for plant availability.

## Acknowledgments

This work was financially supported by the Office National de la Chasse et de la Faune Sauvage. L’Office National des Forêts (ONF) provided funding for vegetation surveys through the agreement “convention cadre Cemagref/ONF”. We are grateful to Daniel Delorme, Pascal Normant, Gérald Goujon, Jean-Pierre Hamard, Olivier Widmer, Thierry Chevrier, Antoine Mételli, Philippe Ballon and the many field operators for field assistance and vegetation surveys. We would like to thank Maryline Pellerin and Eric Marboutin for constructive comments on a previous draft of this manuscript.

## Supplementary material 1: spatial-autocorrelation

Our sampling design of vegetation plots is a 300×300m grid overlaid on the Territoire d’Étude et d’Expérimentation of Trois-Fontaines. Neighbouring sampling plots may therefore be more alike in plant availability and level of browsing, than between more distant ones. Statistically speaking, spatial-autocorrelation might occur in our data, breaking the assumption of independence of observations of generalized linear models (GLM). Several methodological options were developed to account for spatial-autocorrelation in statistical models (see Dormann et al. 2007 for a review).

In our case, spatial-autocorrelation was a noise parameter and of no biological relevance. We therefore chose to account for spatial-autocorrelation in our data by fitting a Generalized Additive Model (GAM, Hastie & Tibshirani 1987), entering geographic coordinates in the model as smoothed, interactive terms (see also Woods 2017). We did not estimated spatial auto-correlation, nor did we modeled it explicitly. Ultimately, we wanted to compare the estimated coefficients for the effect of density of roe deer, plant availability and the first order interaction on browsing, with and without accounting for spatial-autocorrelation (GAM vs GLM). This was to assess the robustness of our results to the issue of spatial auto-correlation.

We shall recall the structure of the best GLM describing our data:

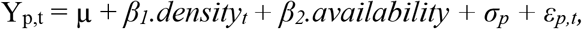

our model hence included the effect of roe deer density, plant availability, a random effect of the sample plot id (*σ_p_*), and an error term (*ε_p,t_*).

As a first step, we quantified the spatial autocorrelation by calculating Moran’s I statistics applied on the GML residuals (population level residuals). Here we used the *gearymoran* function implemented in the R package *ade4*. We found little evidence of autocorrelation in the model’s residuals of roe deer browsing at Trois-Fontaines. Applied to our data, Moran’s *I* = −6.413232e-05 (sd = 0.4894084; *p* = 0.267) was not statistically significant. We hence expect that accounting for spatial auto-correlation or not in our statistical model should have very limited effect, if any.

In a second step, we built a similar GAM with the same fixed and random structure as the selected GLM, along with the smoothing splines for X and Y coordinates and the interaction between X and Y. This modeling approach allows to capture rather complex spatial structure in the data, while minimizing the number of parameters needed for its description. We used the MGCV R package to fit this model to our data (Woods 2011).

**Fig. S1:**
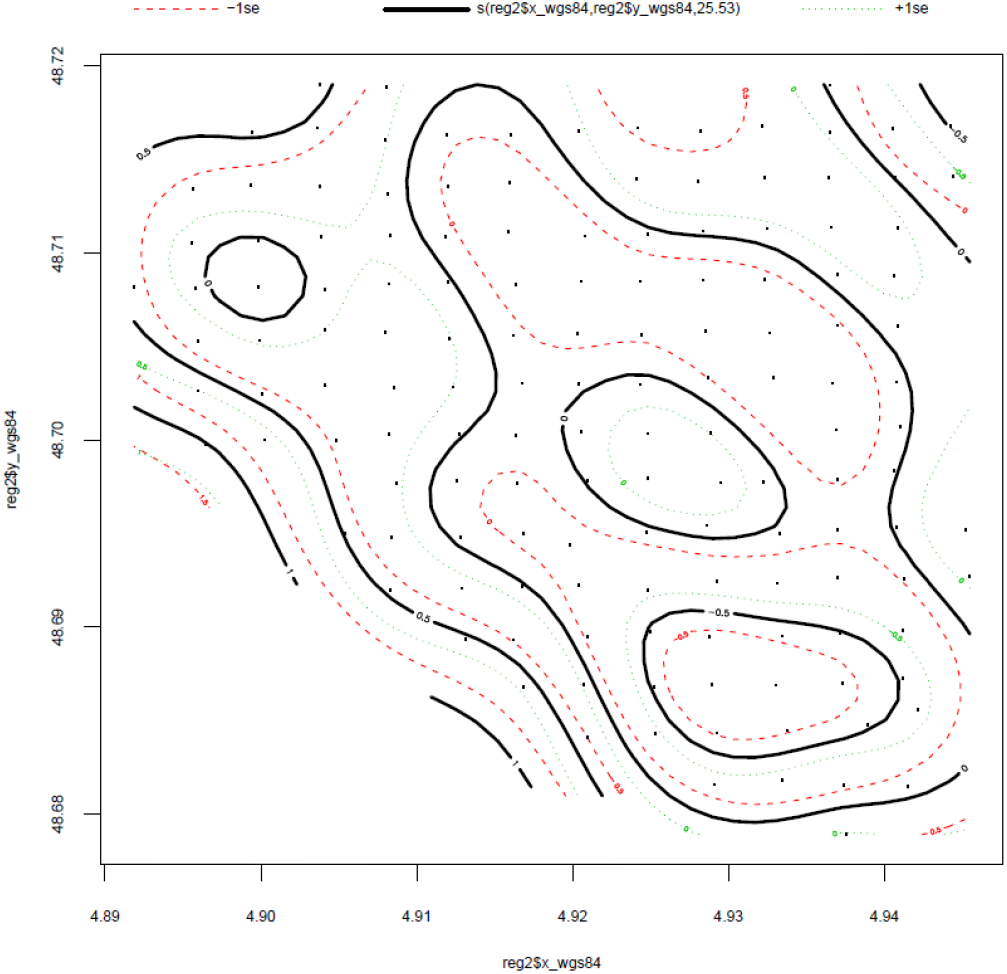
Spatial structure in the browsing of woody plants by roe deer at Trois-Fontaines (France). Small black dots are the location of each sampling plot of the vegetation. Contour (black lines) are isoclines representing the space for which deviation of browsing from the mean value is the same. Coloured dotted lines are the 95% confidence limits of the associated isocline.

We evidence a marked spatial structure in the browsing of woody plants by roe deer at Trois-Fontaines (Fig. S1). Coefficient associated to smoothing splines for coordinates are all significant according to the GAM fit:

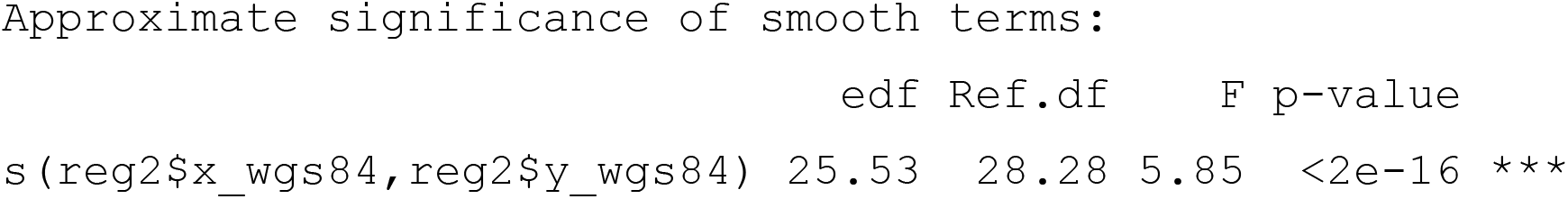

Despite this marked spatial structure in our data, the effects of roe deer density and availability were remarkably similar between the GLM and GAM estimates, i.e. when accounting for spatial auto-correlation or not. The mean relative difference calculated over the 5 parameters was 2%, a value we consider as negligible.

**Fig. S2:**
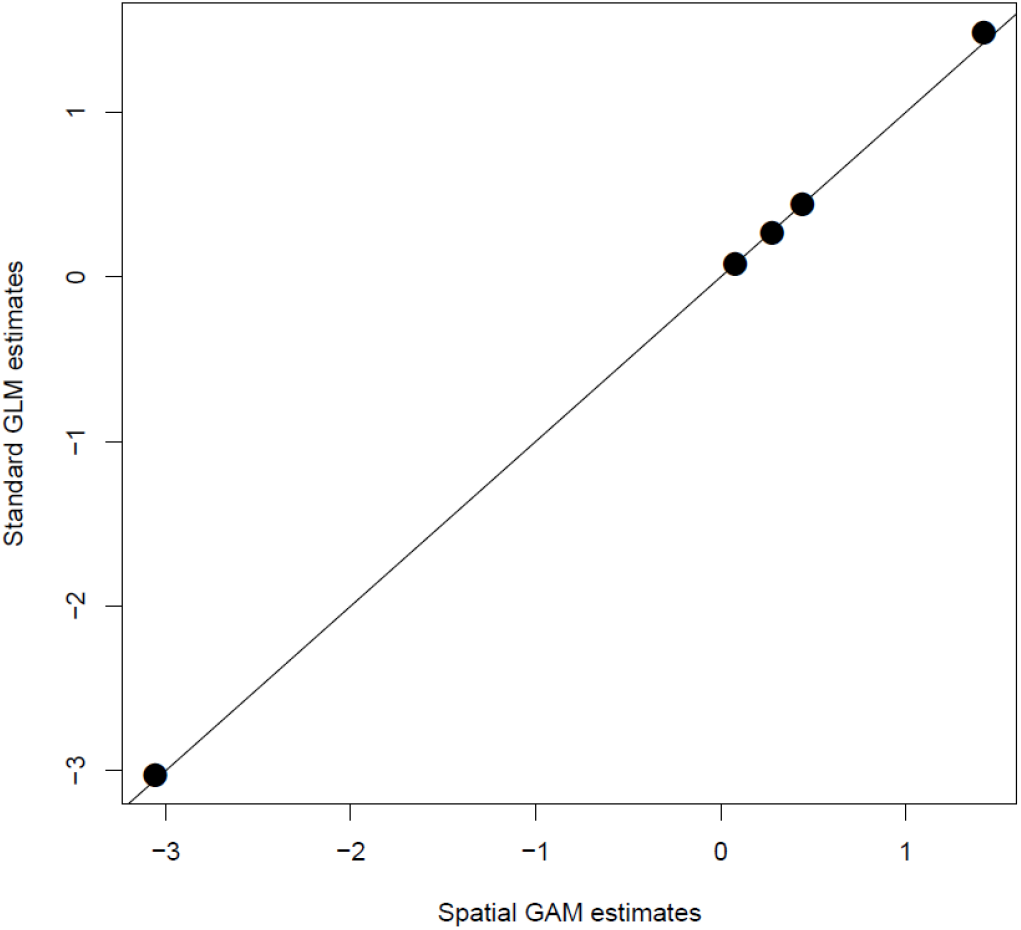
Coefficients for the effects of roe deer density and plant availability on the observed browsing at Trois-Fontaines, France, as estimated from a standard Generalized Linear model not accounting for spatial auto-correlation in the data, and from a Generalized Additive Model, entering plot coordinates as smoothers, hence accounting for the spatial autocorrelation. The plot clear shows that accounting for spatial auto-correlation was unnecessary in our study.

We conclude our results are very robust with regards to spatial auto-correlation issue. We therefore present the analyses with standard GLMs in the main text for the sake of simplicity.

